# Molybdate to prevent the formation of sulfide during the process of biogas production

**DOI:** 10.1101/2019.12.12.874164

**Authors:** Pietro Tenti, Samuele Roman, Nicola Storelli

## Abstract

The process of anaerobic digestion producing biogas is an eco-friendly energy source that promotes recycling from waste biomass such as food chain residues, wet waste, wastewater. In this study, we focused on the problem of the sulfide (H2S) produced by the sulfate-reducing bacteria (SRB) in the presence of sulfate residues. This byproduct is dangerous for human health and an issue due to the highly corrosive effect on metallic components. To this purpose, the Molybdate, a sulfate analog, known in the literature to inhibit SRBs by blocking the first enzyme of the metabolic pathway of anaerobic respiration, was applied. The experiments carried out showed that a concentration of 0.3 mM of molybdate was enough to inhibit the SRB in a complex environment of the anaerobically digested sludge (ADS) took from a real biogas producing bioreactor. During the study, we observed the importance of the sulfate concentration sulfate in the system. Indeed, the production of sulfide was stopped only under the threshold ratio value of 1:10 (molybdate: sulfate). In the short term, the addition of molybdate did not alter the production and quality (% of methane) of the biogas, nor the anaerobic microbial community, including SRB population itself.

## INTRODUCTION

Developed countries are starting to develop the concept of a “circular economy,” in order to reduce waste and pollution in order to guaranty a decent future of the planet. The main aims of this model are the recycling of goods and waste, the increase in the production processes quality, and the prolonge of the products’ life. The process of anaerobic digestion producing biogas can produce energy from organic wastes and is an environment-friendly (compared to thermo-physical processes) and a valid alternative to fossil fuel ^1,2^. This biological process involves different communities of anaerobic microorganism responsible for degrading organic matter and producing biogas as final product (approximately 60% methane and 40% CO_2_), which is used to produce electrical energy in different ways ^3,4^. In anaerobic bioreactors, the presence of sulfur compounds (e.g., protein, sulfate, thiosulfate, sulfite) leads to the formation of highly toxic, corrosive, and malodorous hydrogen sulfide (H_2_S). Sulfate-reducing bacteria (SRB) reduce sulfate (SO_4_^2-^) to sulfide by transferring electrons obtained by the degradation of organic matter such as lactate and acetate. The presence of SRB decreases the final biogas production by competing with methanogen for acetate and by generating toxic H_2_S ^5–8^. Values higher than 500 ppmv of sulfide in the biogas, produce a dangerous corrosive effect, significantly reducing the lifetime of pipes and other metallic hardware ^9^. The concentration of H_2_S in biogas was reported to usually range from 50 to 5,000 ppmv with peaks of up to 20,000 ppmv (2% v/v) in some cases ^10^. Moreover, in the presence of an excess of sulfate, SRB outcompetes with methanogens for the common substrates such as hydrogen and acetate ^11,12^. Owing to the higher affinity and lower threshold values for hydrogen, methanogens are easily and rapidly out-competed by SRB ^7^.

At the present day, the removal of the sulfide from the biogas is an on-going issue without a clear dominant solution ^9,10,13,14^. Three methods are mostly used to solve the problem: 1) chemical precipitation/oxidation frequently using iron chloride (FeCl_3_) and gaseous oxygen (O_2_), 2) ionic absorption using activated carbon or zeolite and 3) biological (chemotrophic) degradation using microaerobic colorless sulfur bacteria from the genus of *Thiobacillus* sp. Each method removes the sulfide from the biogas, but there are also negatives points to take into account, such as products costs, use of oxygen in an anaerobic process, instability through fluctuating sulfide concentrations, environmental problems given by the production of toxic products or by-products, and considerable disposal costs. Additionally, all cited methods remove the H_2_S after its formation and are therefore not entirely effective against the damage of metallic components triggering high maintenance cost, estimated to be US$2.5 trillion ^15^.

Past studies already investigated the possibility of inhibiting SRB using different strategies. For example, the application of nitrate ions (NO_3_^-^) stimulates the growth of nitrate-reducing bacteria resulting in competition with the SRB for the available carbon source in the environment ^16–19^. This method is an attractive solution because it is cheap, relatively non-toxic, and easy to apply in big reservoirs. However, nitrate-reducing bacteria compete as well with methanogens decreasing the final biogas quality.

The molybdate ion (MoO_4_^2-^), similar to other analogs of the sulfate (chromate, tungstate, and selenite), inhibits sulfate reduction of SRB. Molybdate enters cells via a sulfate transport system and interferes with the formation of adenosine phosphosulfate (APS). The formation of adenosine phospho-molybdate in the cell inhibits the following reduction to (bi)sulfite by APS reductase (AprBA) and stop the respiration pathway ^20,21^. The specific inhibitory action of the molybdate showed a reduction of sulfide concentration when applied to microbial sulfate reduction processes ^22–24^.

In this paper, we evaluate the inhibitory effect of the molybdate in the process of anaerobic digestion at the laboratory scale.

## MATERIALS AND METHODS

### Microbial culture

The anaerobically digested sludge (ADS), used as active biomass in all experiments, was taken from two bioreactors of 4000 m^3^ routinely producing biogas (Gordola wastewater treatment plant WWTP; Consorzio depurazione acque del Verbano, Switzerland). Inhibition tests were carried out in 125 ml sealed glass bottles inoculated with 60 ml of ADS, diluted 1:4 with tap water, and flushed with an anaerobic gas mixture (10% H_2_, 10% CO_2_ and 80% N_2_) to remove oxygen. The 1:4 dilution ratio was defined in order to allow the use of syringe and needles in the process of inflow and outflow of the samples.

In the first experiment (evaluation of the molybdate inhibitory effect) four different concentrations of sodium molybdate (Na2MoO4; 1.0 mM, 0.6 mM, 0.3 mM, 0.1 mM) was added to three bottles (triplicate), from a sterile stock solution of 1.0 M in demineralized water. All samples were incubated at 37°C with a constant daily inflow/outflow of 2.0 ml until the end of the experiment (after 63 days) in order to simulate a bioreactor activity. Sterile sodium acetate (NaC_2_H_3_O_2_; 2 ml; 33.3 mM) from a sterile stock solution of 1.0 M in demineralized water was used as a daily inflow. The use of acetate was chosen to stabilize the methane-producing microorganisms (acetoclastic methanogens). The outflow of 2.0 ml from the bottles was expected to reduce the molybdate concentration in the system of 33.0 µM, 20.0 µM, 9.9 µM, and 3.3 µM per day, respectively. After16th day of incubation, one single injection of 1.6 g/l of sterile anaerobic LB rich medium (yeast extract and bacto-peptone) was added in order to provide enough energy to the microbial community.

In the second experiment (influence of the sulfate concentration), the inhibition activity provided by a determined concentration of 0.8 mM molybdate was evaluated in the presence of different concentrations of sulfate. All the 125 ml sealed glass bottles (triplicates), except for the control bottle (triplicate), were prepared with an initial concentration of 0.8 mM of sodium molybdate (Na_2_MoO_4_), from a sterile stock solution of 1.0 M in demineralized water. Sodium sulfate (Na_2_SO_4_) was then added from a sterile stock solution of 1.0 M in demineralized water, based on the concentration of sulfate already present in the sample resulting in different ratios with the molybdate of 1:4, 1:10, 1:20, 1:40, 1:60. All samples were incubated at 37°C in batch mode (no daily inflow and outflow as in the previous experiment). After 24 hours of acclimation, 1.6 g/l of sterile anaerobic LB rich medium (yeast extract and bacto-peptone) was added once to every bottle in order to provide enough energy to the microbial community.

### Laboratory-scale bioreactors and feeding procedure

Two equal lab-scale Continuous-flow Stirred-Tank Reactors (CSTR) of 3.0 liters were filled with 1.0 liter of ADS from the WWTP, flushed with an anaerobic gas mixture (10% H_2_, 10% CO_2_ and 80% N_2_) to remove oxygen and incubated at 37°C with 1.0 hour interval of magnetic stirring at 150 rpm every 2.0 hours of rest. Both lab-scale CSTRs were fed daily with 60.0 ml of various organic waste provided by Gordola WWTP for hydraulic retention time (HRT) of approx. 16.5 days were intending to simulate the one applied to the full-scale 4000 m^3^ anaerobic digester. The only difference between the two CSTRs was the method used to reduce/remove the sulfide. In one CSTR, a concentration of around 0.5 mM of molybdate was maintain along with the experiment, while in the second CSTR, iron chloride (FeCl_3_) was added to oxidize sulfide once measured.

In the first two weeks, both CSTRs were fed using a 1:4 diluted sewage sludge (COD of about 1.0 g/l) corresponding to an organic loading rate (OLR) of 0.021 g/l day. In the third and fourth week, undiluted sewage sludge (COD of about 4.2 g/l) was inoculated, corresponding an OLD of 0.084 g/l day. From the fifth to the ninth week, cheese whey (CW) was added to the sewage sludge in a 1:4 ratio, according to the procedure applied at the WWTP. The first three weeks was used a CW with low-fat content (COD of about 50.0 g/l) and the remaining two weeks with a CW with high-fat content (COD of about 80.0 g/l), thus providing respectively an OLR of 1.0 and 1.6 g/l day. In the tenth week, homogenized waste from restaurants and canteens (RCW) was added to the sewage sludge in a 1:4 ratio, with an OLR of 3.6 g/l day. In the eleventh and twelfth weeks, concentrated industrial fermentation media, with a high concentration of amino acids (especially methionine) and sugars, were added to the sewage sludge in a 1:4 ratio, with an OLR of 4.0 g/l day. In the last week, both CSTRs were inoculated again with 1:4 sewage sludge as in the third and fourth weeks to reduce the biogas production and stop the experiment.

### Chemical measurement

Chemical oxygen demand (COD), alkalinity, ammonium, sulfate, and sulfide were measured from the supernatant after centrifugation (10 min at 5000 rpm) using specific colorimetric chemical kits provided by Merck AG (Zug, CH), according to the manufacturer’s instructions. The concentration of acetate was measured in the same manner but using the r-Biopharm (Darmstadt, Germany) specific acetic acid kit according to the manufacturer’s instructions.

The methane concentration in the biogas was determined by gas chromatographic separation and subsequent determination with a flame detector (FID) using the Varian 450-GC gas chromatograph (Varian, USA) coupled to a hydrogen generator. Samples were analyzed at room temperature for a total of ten injections per single test sample. The chromatographic separation column used was a Guard Column 5m x 0.53, i.d. deactivated tubing. The flame detector uses hydrogen, which is mixed with air and nitrogen, the eluent of the column, burning in the small nozzle inside a cylindrical electrode; a potential of 100 V was applied between the nozzle and the electrode. When a sample containing carbon was burned, the electron/ion pairs formed are detected. The instrument releases a result in μV.sec representing the area subtended to a peak after an injection from which the μg/L of methane produced by both CSTRs were automatically determined by inserting the μL of sample injected during the analyzes. The percentage of methane produced from 1 liter of culture was determined according to the following formulas: first, it was calculated the total moles present in two liters of reactor atmosphere with PV = nRT with P = internal pressure at the reactor, n = 0.0821, T = 310K. Subsequently, using the CH_4_ PM, the moles of CH_4_ produced in two liters are determined. Finally, by dividing the moles of CH_4_ for the total moles and multiplying by 100 to obtain the percentage present in the biogas.

### Flow cytometry counts

The coenzyme F420 involved in methanogenesis causes an intense auto-fluorescence of cells under excitation by shortwave UV light (max Abs 420nm). This auto-fluorescence is a diagnostic feature and can be used to check the cultures of methanogens by flow cytometry. BD Accuri C6 cytometer (Becton Dickinson, San José, CA, USA) equipped with two lasers (488 nm, 680 nm), two scatter detectors, and four fluorescence detectors (laser 488nm: FL1 = 533/30, FL2 = 585/40, FL3 = 670; laser 640 nm: FL4 = 670) has been used for this purpose. Flow cytometer parameters have been used for events characterization: forward scatter (FSC), which is often correlated to particle size, 90° light scatter (SSC), which is considered to be related to the size and internal granularity of the particles. Thresholds have been applied first to forward scatter (FSC-H 10’000), for the exclusion of debris and abiotic particles, and subsequently to the FL1 filter for the detection of natural fluorescence of coenzyme F420.

### Fluorescent *in situ* hybridization (FISH)

The total SRB and the acetoclastic methanogens of the genus *Methanosarcina* have been identified and counted with species-specific Cy3-labeled oligonucleotides SRB385 (CGGCGTCGCTGCGTCAGG) ^25^ and the MS821 (CGCCATGCCTGACCTAGCGAGC) ^26^ respectively, with 2 and 5 µl aliquots of paraformaldehyde-fixed samples (n = 3) spotted onto gelatin-coated slides [0.1% gelatin, 0.01% KCr(SO_4_)_2_] ^27^. Hybridizations were performed, as described in previous studies. Slides have been treated with Citifluor AF1 (Citifluor Ltd., London, UK) and examined by epifluorescence microscopy using filter sets F31 (AHF Analysentechnik, Tübingen, Germany; D360/40, 400DCLP, and D460/50 for DAPI) and F41 (AHF Analysentechnik; HQ535/50, Q565LP, and HQ610/75 for Cy3). Microorganisms were counted at a 1’000 fold magnification in 40 fields of 0.01 mm^2^ each.

## RESULTS

### Evaluation of the molybdate inhibitory effect

The inhibition effect of the molybdate on the SRB respiratory chain was already confirmed using pure cultures of *Desulfovibrio* ^20^. However, the efficacy of this process in the production of biogas through anaerobic digestion is still not proved yet. The hydrogen sulfide (H_2_S) production of active anaerobically digested sludge (ADS) samples coming from the bioreactor of the Gordola WWTP was evaluated with the addition of different concentrations of molybdate (Figure 1A). During the first phase of 15 days, all the samples were fed daily with only a sterile anaerobic solution containing acetate (33.3 mM), in order to remove any trace of oxygen, unknown remaining organic energy, and to stabilize the acetoclastic methanogens. During this period in all samples, included negative control without molybdate, no trace of sulfide and no internal pressure due to the biogas production was recorded. Flow cytometry measurement also suggested low metabolic activity of the anaerobic microbial community (Figure 1B). Indeed, the methanogens decreased from 6.1 to 3.2 10^6^ cells/ml, whereas the total counts decreased accordingly from 8.6 to 5.1 10^7^ events/ml. After 15 days the energetically content of the system was almost zero (COD = 3’000 mg/l).

**Figure 1:**
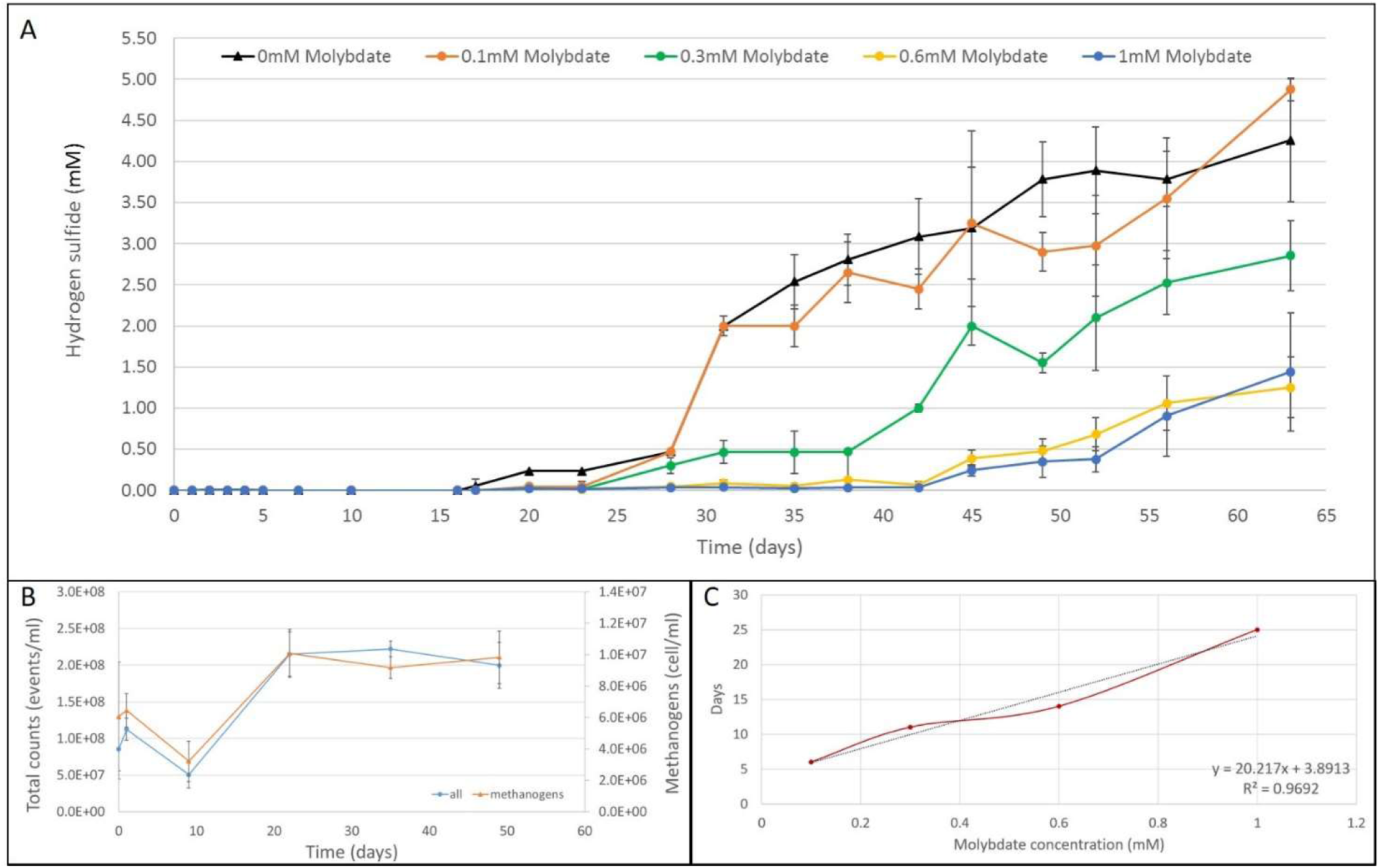
(A) Hydrogen sulfide (H_2_S) production (mM) recorded in 125 ml sealed bottles (triplicate) with different concentrations of molybdate. The negative control without molybdate is the black line (black triangle), the blue line (blue dot), the yellow (yellow dot), the green (green dot) and the orange (orange dot) lines correspond to the starting molybdate concentration of 1.0 mM, 0.6 mM, 0.3 mM, and 0.1 mM, respectively. (B) Flow cytometry counts of methanogens and all microorganisms in the ADS. The blue dots correspond to the overall events counted per ml, while the orange triangle to the auto-florescent methanogens per ml. Every point is a mean value of the counts for the 5-incubation conditions due to the minimal difference in terms of microbial community composition. (C) Length of inhibition (days) in the function of the starting concentration of molybdate present in the sample (1.0 mM, 0.6 mM, 0.3 mM, and 0.1 mM).

**Figure 2:**
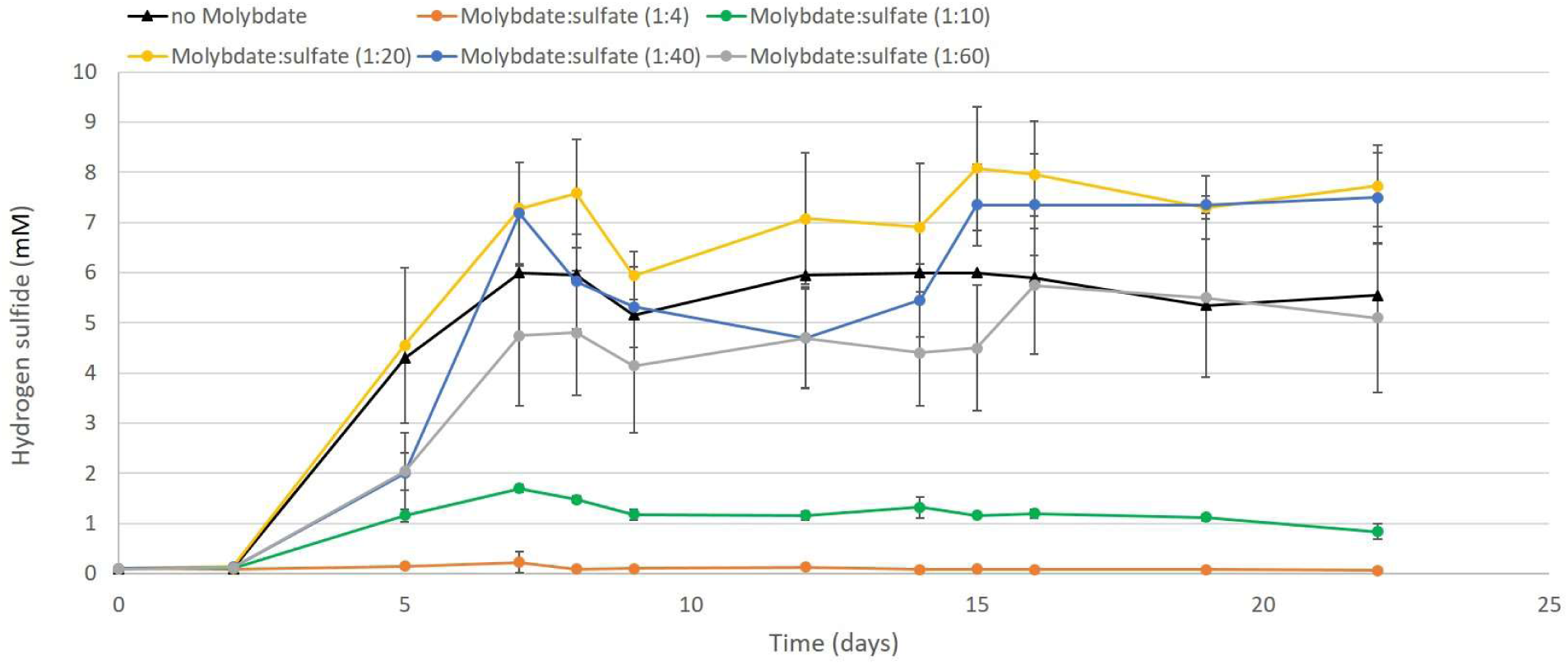
Hydrogen sulfide (H_2_S) production (mM) recorded in 125 ml sealed bottles (triplicate) with different ratios of molybdate (MoO_4_^2-^) and sulfate (SO_4_^2-^). The negative control without molybdate is represented by the black line (black triangle), ratio 1:4 orange-, ratio 1:10 green-, ratio 1:20 yellow-, ratio 1:40 blue- and ratio 1:60 grey-line.

The inhibition effect of the different concentrations of molybdate was finally tested after the addition of LB media in every sample (day 16) increasing of 10-fold the energy to a COD value of 35’000 mg/l. The following day (day 17), the concentration of sulfide was finally of 0.05 mM in the negative control bottles (Figure 1A, black triangle). On day 20, the concentration of sulfide increased to 0.23 mM and appeared in the bottles with minor concentration of molybdate of 0.1 mM (Figure 1, orange dots). After 12 days (day 28), traces of sulfide was also measured in the bottles with 0.3 mM and 0.6 mM of molybdate, with values of 0.3 mM and 0.05 mM, respectively. Finally, after 27 days (day 43), the H_2_S increased in the last sample containing 1.0 mM of molybdate. At the end of the experiment (day 63), the final concentrations of sulfide in the examined samples were 4.26 for the negative control and 4.88, 2.86, 1.25, and 1.44 mM, for bottles with starting concentration of molybdate of 0.1 mM, 0.3 mM, 0.6 mM, and 1.0 mM, respectively. After the increase of the energy in the bottles (day 16), flow cytometry data showed a rapid increase of the total microbial community and methanogen cells that remained stable until the end of the experiment (Figure 1B).

At the end of the experiment, the time of inhibition in the function of the starting concentration of molybdate was calculated, taking into account the washout effect due to the inflow/outflow of sludge (Figure 1C). The result clearly showed a linear relationship between the concentration of molybdate and the length of the inhibition mechanism (R^2^=0.9692).

### Influence of the sulfate concentration

Molybdate (MoO_4_^-2^) is a structural analog of sulfate (SO_4_^2-^) with a high affinity for the PAPS synthetase enzyme, so its presence inhibits the respiration of SRB ^20^. Five different concentrations of sulfate were analyzed in the presence of a fixed amount of molybdate (0.8 mM) to investigate the ionic competitory effect and a possible subsequent reduction of inhibition effect. In this second experiment, the stabilization period was reduced to one single day because of the insufficient level of energy in the starting 1:4 diluted ADS. After 24 hours all cultures were fed only once with LB media corresponding to a starting COD value of approx.. 40’000 mg/L.

Hydrogen sulfide appeared after 48 h (day 2) of incubation in control and samples with molybdate: sulfate ratios higher than 1:20 (Figure 3, graphs yellow 1:20, blue 1:40, and grey 1:60). Samples with 1:10 molybdate: sulfate ratio (Figure 3, green graph) also showed the presence of a small quantity of H_2_S after 48h (day 2) increasing until a maximal value of 1.7 mM after seven days, but then slightly decreasing and stabilizing at approx. 1.2 mM until the end of the experiment. No production of H_2_S was observed for samples with 1:4 molybdate: sulfate ratio during the experiment (22 days); the concentration of H_2_S remained the same at approx. 0.1 mM (Figure 3, orange graph). In this case, the sulfate concentration was not sufficient to compete with the inhibition effect of molybdate on the SRB community.

**Figure 3:**
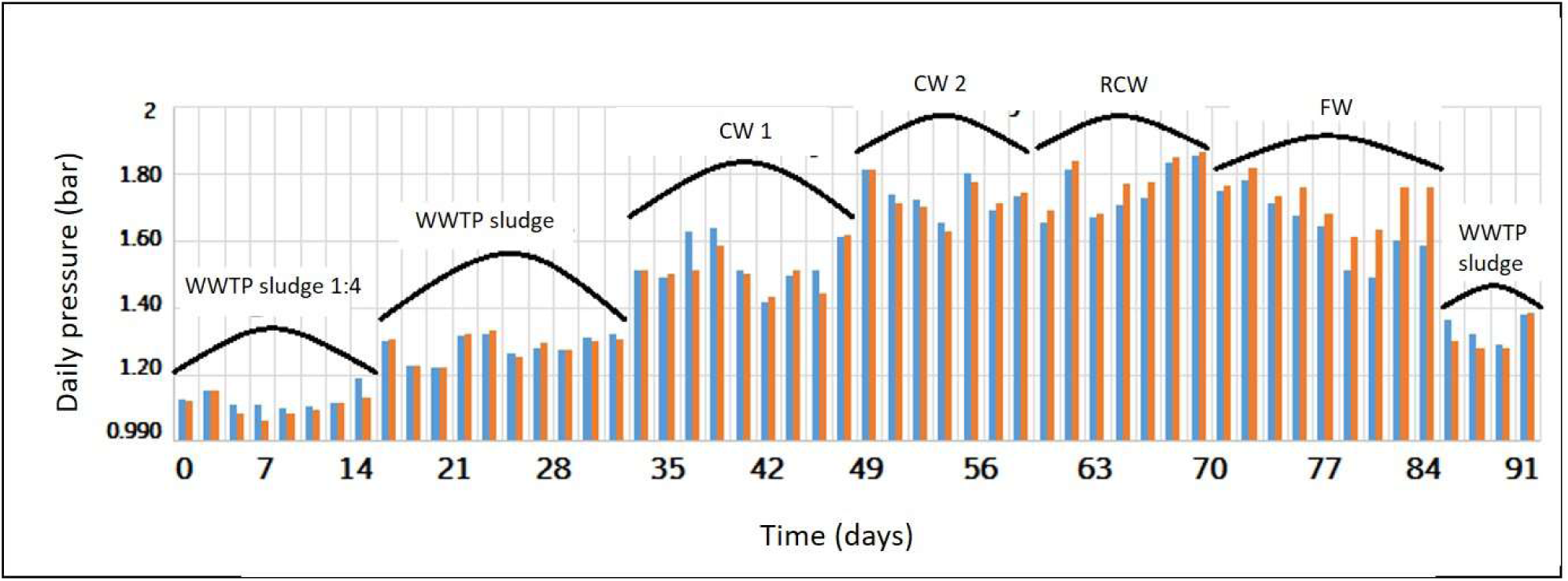
Daily pressure (bar) generated by both CSTRs with or without molybdate in relationship with the organic waste used as energy source. The orange bars are measured from the bioreactor containing molybdate and the blue bars from the control without molybdate. Every 2-3 weeks, the organic waste added change, increasing the OLR.

The specific SRB counts by FISH at the end of the three weeks of the experiment did not show variation in the number of SRB cells staying at approx. 4.5 10^8^ cells/ml in every sample. This data suggest that SRBs were inhibited but not killed by molybdate.

### Laboratory-scale biogas production using molybdate

The biogas production of two parallel 3.0 liters CSTR was evaluated by the use of two different methods to prevent or neutralize the sulfide. In the first bioreactor, a minimal concentration of 0.5 mM of molybdate were kept almost constant during the experiment period (92 days). In the second bioreactor, the level of sulfide was reduced using iron chloride (FeCl_3_), which acts as a scavenger, oxidizing sulfide to pyrite (FeS_2_). Both CSTRs were set-up using the real biogas bioreactor of the WWTP of Gordola as a model system (see Materials and methods).

The first substrate was the sewage sludge produced by the process of wastewater purification; moreover, the WWTP of Gordola regularly included various organic wastes such as cheese whey (CW), canteen and restaurant waste (CRW), and fermentation waste (FW) from industrial microorganism production. The evaluation started with a dilution 1:4 with water of the WWTP sewage sludge and after two weeks (14 days) without dilution for further three weeks (35 days) in order to stabilize the process of anaerobic digestion. After the first stabilization period every two weeks, the OLR was increased continuously by adding to the WWTP sludge approx. 25% of various organic wastes, starting from the less energetically CW1 (COD around 100’000mg/l) until the most one FW (COD around 400’000 mg/l). In the final period (10 days) of the experiment, both bioreactors were fed only with WWTP sludge to reduce biogas production. The amount of daily biogas produced correlated well with the OLR; high values resulted from high energetically organic waste (Figure 3; blue and orange bars, for control and molybdate respectively).

The first measured value of H_2_S appeared after 72 days in both bioreactors respectively 0.048 mM in the first CSTR with molybdate, and 0.092 mM in the second CSTR without molybdate (Figure 4). The concentration of the sulfate (SO_4_^2-^) increased only in the bioreactor with the molybdate (Figure 4, yellow diamonds). In fact, due to the inhibition of SRB, the sulfate was not reduced in sulfide and accumulated in the bioreactor. Therefore, it was necessary to increase the concentration of molybdate to 1.2 mM to maintain the ratio with the sulfate under 1:10 (showed in the previous experiment). The increase in molybdate concentration stopped sulfide production for approximately one week (day 81), before the concentration of sulfate increased again over the 1:10 ratio (Figure 4, red triangle). After a while, the experiment was stopped, and energetically poor WWTP sludge was added to reduce the production of biogas (Figure 3).

**Figure 4:**
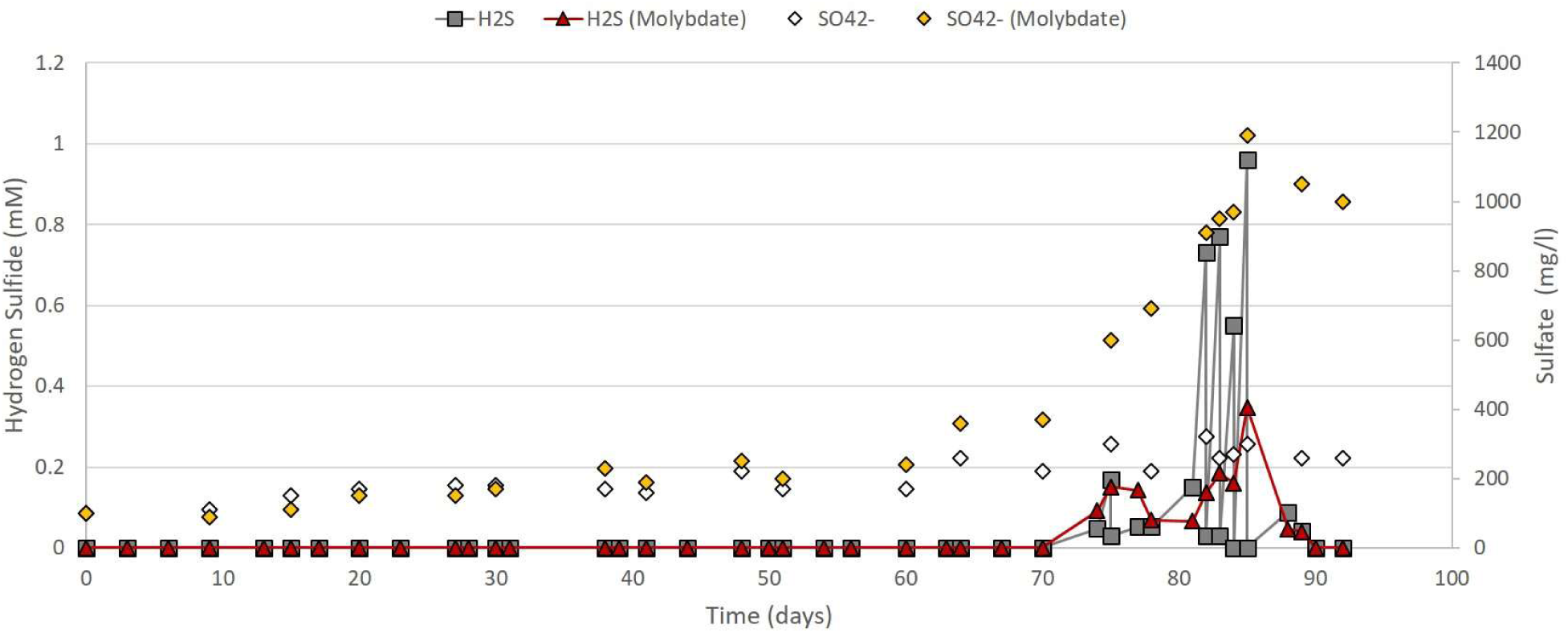
Values of the hydrogen sulfide (H_2_S) and sulfate (SO_4_^2-^) produced along the experiment from both CSTRs. In the first CSTR with molybdate, the red triangle graph corresponded to the H_2_S and the yellow diamonds to the SO_4_^2-^. In the second CSRT without molybdate, the grey square graph corresponded to the H_2_S and the white diamonds to the SO_4_^2-^.

During the experiment, the concentration of alkalinity (HCO_3_^-^), of ammonium (NH_4_^+^), and methane in the biogas (%CH_4_), were regularly monitored (Figure 5), and all showed similar values for both CSTRs. The percentage of methane in the biogas increased together with the content of energy in the added substrate (Figure 5, red, and white bars), showing a high 93% on day 88 of the measurement. The increase of methane also increased the alkalinity reaching maximal values of approx. 9’000 mg/l at day 83. Whereas, the concentration of ammonium remained constant around 500-600 mg/L along with the experiment.

**Figure 5:**
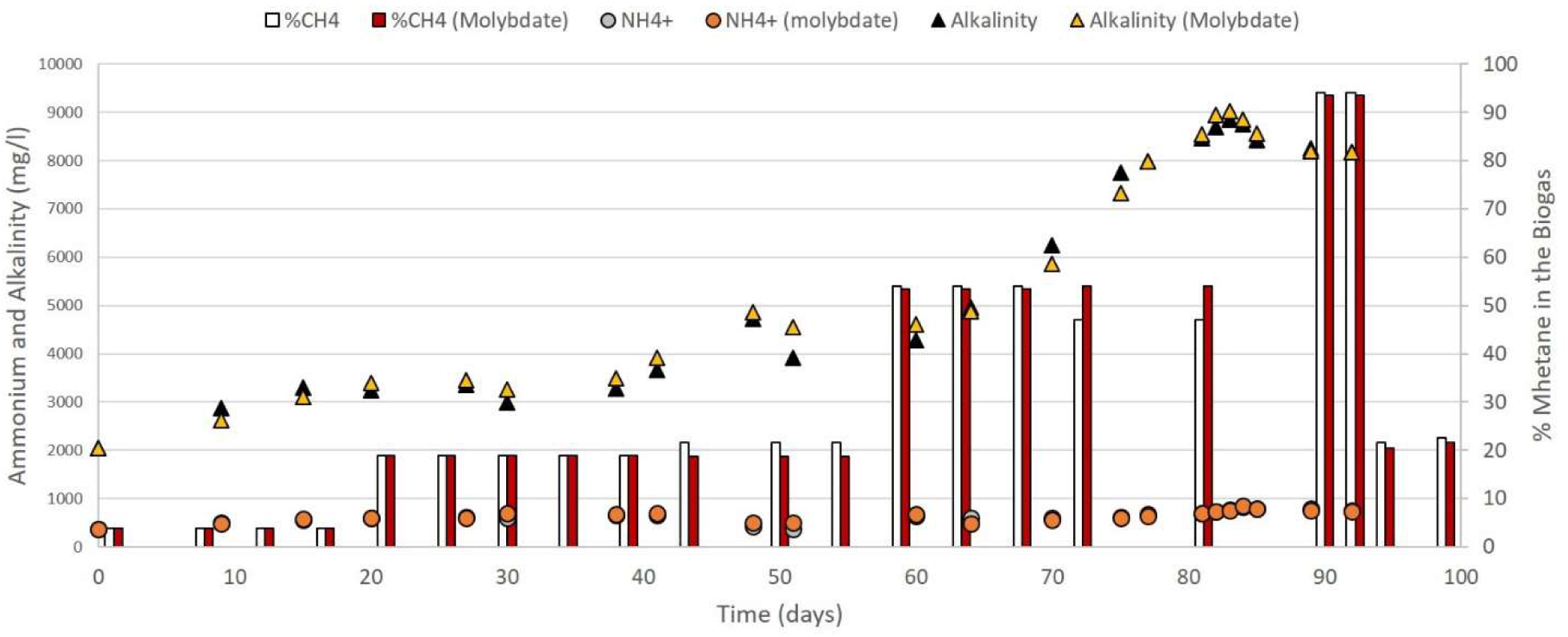
The concentration of alkalinity (HCO_3_^-^), ammonium (NH_4_^+^), and methane in percentage (CH_4_) were monitored continuously during the experiment. Values of alkalinity are presented with a yellow and black triangle for the CSTR with and without molybdate, respectively. Amounts of ammonium are shown with an orange and gray circle for the CSTR with and without molybdate, respectively. The percentage of methane in the biogas is displayed with red and white bars for the CSTR with and without molybdate, respectively.

At the end of the period of measurement, the microbial community of both lab-scale bioreactors (with and without molybdate) was evaluated. Two specific fluorescent probes for all SRB and the acetoclastic methanogens of the genus *Methanosarcina* were used. The concentration of total cells, SRBs, and methanogen microorganisms were almost the same in both tested bioreactors (Table 1).

**Table 1.**
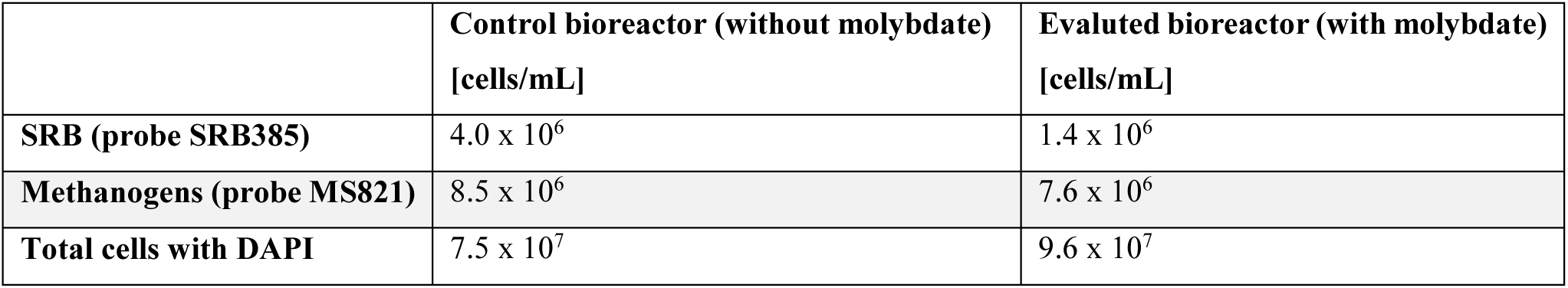
Total number of SRB (probe SRB385), *Methanosarcina* (probe MS821) and all microorganisms (DAPI) characterized by FISH.

## DISCUSSION

One of the main problems in the production of biogas through anaerobic digestion is the production of highly toxic, corrosive, and malodorous sulfide. In this study, an inhibition effect of molybdate on the SRBs respiratory activity is observed starting from a concentration of 0.3 mM up to a concentration of 1.2 mM without affecting the process of anaerobic digestion and the resulting biogas production. The efficacy of the inhibition is strongly correlated with the sulfate concentration; in fact, ratios higher than 1:10 molybdate: sulfate failed to stop the production of sulfide. The batch experiment designed to evaluate the influence of sulfate on the molybdate inhibition suggested that SRBs were not able to metabolize molybdate during 20 days (Figure 2 green and orange graph). The concentration of SRBs in the samples influenced the efficiency of the inhibition; in fact, inhibition of SRBs was observed at molybdate doses of 0.5 mM or higher in SRB enriched biomass, whereas similar inhibition is observed at the relatively lower dose of 0.25 mM in SRB deficient biomass ^22^. The inhibitory effect of molybdate in pure cultures of SRB was even higher, with a minimum inhibitory concentration of 0.08 mM ^23^. The threshold of inhibition was also related to the sulfate concentration in pure cultures; for example, in incubation media containing 20.0 mM sulfate, the production of sulfide was stopped only above two mM molybdate, corresponding to an adequate ratio of molybdate:sulfate of 1:10 similar to our observation ^24^. To reduce the concentration of sulfide in the biogas, Peu et al. suggested the importance of also keeping the carbon: the sulfur ratio of the organic waste under 40 ^28^. All these data indicated the importance of choosing the correct feedstock in power plants aiming to increase the quantity and quality of the produced biogas. Not only the feedstock but also the pH seemed very important in influencing the process of anaerobic digestion. Indeed, the rise of the initial sludge pH from 6.5 to 8.0 inhibited the competition of SRBs with methanogens, and thus promoted the growth of methanogens and the biogas production ^29^. Nowadays, there are many physical-chemical and biotechnological technologies for the removal of H_2_S from biogas ^13,30,31^. Given the wide choice of possibilities, the concentration of H_2_S in the biogas is a fundamental parameter for the choice of the best desulfurization technology. A precise estimation of the H_2_S concentration is therefore of fundamental importance to decide whether or not the installation of a desulfurization technology is necessary (concentration up 500 ppmv) ^32^.

The maximal concentration of sulfide reached values up to 5.1 mM (163 mg/l). From the literature, it is known that above a specific concentration, sulfide inhibits first the methanogens, and then all other bacterial communities included the SRBs themselves ^33^. Similarly, high concentrations of molybdate up to 2.5 mM were negative for the methanogen activity and the consequent production of biogas ^22,34^. Moreover, with a molybdate concentration below 1.2 mM, the inhibition effect on SRBs was reversible. Indeed, FISH counts resulted in a similar number of total SRB around 5.0 10^8^ cells/ml in all samples, with or without molybdate, at the end of every experiment. SRBs FISH cell detections were supposed to be positive even with reduced cell activity by the presence of a minimal amount of ribosomal RNA, indicating metabolic activity and, therefore, protein synthesis ^35^. Also, cell counts obtained with FCM supported the same interpretation, namely that low concentration of molybdate did not influence the metabolism of the methanogens and, more generally, of the microbial community. The harmless of molybdate was also confirmed in the unvaried levels of biogas production (Figure 3).

From an economic point of view, molybdate is expansive with an average price of 10 USD/Kg, compared with the much cheaper iron chloride. Although classical used methods are cheaper compared to the use of molybdate, they act a posteriori by eliminating the sulfide only once it has been produced; they cannot eliminate the damages caused by the acid overtime on iron parts of the plant. Molybdate, on the other hand, would have a higher initial cost, but by inhibiting sulfide production, it would avoid damaging expensive plant components such as tanks or pipes. Another positive outcome in using molybdate to prevent sulfide production is the accumulation of sulfate in the digested sludge (Figure 4). Sulfate is essential in agriculture production since it is typically a limiting element. On the other hand, the accumulation of molybdate in the ADS could be dangerous for the environment, but this effect is not completely clear at the moment ^36–38^.

This study gave some interesting indication on the high SRB inhibiting ability of the molybdate (less than 0.5 mM) and the importance of the ratio with the sulfate present in the system (less than 1:10). Regardless, further experiments have to be carried out in order to evaluate the effect of molybdate during a prolonged period and in a bigger bioreactor size, on the process of biogas production. During a prolonged study, it would be possible also evaluate a potential toxicity effect of the molybdate on the anaerobic digestion microbial community and especially after disposal of high quantity of ADS. Moreover, it would be interesting to also evaluate the corrosive effect of the sulfide on all iron hardware of the plant and take into account detailed cost evaluation. The use of bigger and more “professional” bioreactors should provide most precise control on the quantity of molybdate needed in function of the ratio with sulfate in the system. At the present day, the remotion of sulfide from biogas plants using molybdate seems not be possible without further experiments due to the high costs compared to state of the art ^9,14^.

## ACKNOWLEDGEMENTS

We want to thank all people of the laboratory of applied microbiology of SUPSI in Bellinzona, especially the director Professor Mauro Tonolla and Dr. Antonella Demarta responsible for the master student. Moreover, we thank also the Professor Flavia Marinelli of the University of Insubria for accepting the role of rapporteur for the thesis work of Pietro Tenti. We want to thank all LMA colleagues, in particular, Samuele Roman, Francesco Danza, and Federica Mauri, for all pleasant moments and discussions.

## AUTHOR CONTRIBUTION

P.T and N.S contributed conception and design of the study; P.T and S.R carried on all practical experiments; P.T and N.S wrote the manuscript, and N.S prepared all figures. All authors contributed to manuscript revision, read and approved the submitted version.

## ETHICAL STATEMENT

This article does not contain any studies with human or animal participants by any of the authors. No specific permits were required for the described laboratory analysis. Pietro Tenti, Samuele Roman, and Nicola Storelli declare that they have no conflict of interest.

